# Evolutionary variance analysis of the Glycoside Hydrolase Family 13: Structural evidence in classification and evolution

**DOI:** 10.1101/201251

**Authors:** Jose Sergio Hleap, Christian Blouin

## Abstract

Glycoside Hydrolase Family 13 (GH13) structures are responsible for the hydrolysis of starch into smaller carbohydrates. They important in industrial applications and evolutionary studies. This family has been thoroughly documented in the the Carbohydrate-Active enZYmes Database (CAZY), and divided into subfamilies based mainly in sequence information. Here we give structural evidence into GH13 classification and evolution using structural information. Here we proposed a novel method that is sensitive enough to identify miss-classifications, or to provide evidence for further partition that can be of interests to bio-engineers and evolutionary biologists. We also introduced a method to explore the relative importance of residues with respect to the overall deformation that it causes to the overall structure in an evolutionary time scale. We found that the GH13 family can be classified into three main structural groups. There is a hierarchical structure within these clusters that can be use to inform other classification schemes. We also found that by using structural information, subtle structural shifts can be identified and that can be missed in sequence/phylogeny-only based classifications. When each structural group is explored, we found that identifying the most structurally variable sites can lead to identification of functionally (both catalytically and structurally) important residues.

## Introduction

Members of the Glycoside Hydrolase Family 13 (GH13) act on *α*-glucoside linkages of starch. Its members catalyse hydrolysis, transglycosylation, condensation, and cyclization [Ben Ali et al., 2006]. This family of proteins is industrially important in the production of ethanol [Bothast and Schlicher, 2005], high-fructose corn syrup [Visuri and Klibanov, 1987], and other oligosaccharides industrial production. They are also used in the textiles, paper, and detergent industries [Kirk et al., 2002, Gupta et al., 2003]. Biologically, it is also of interest since all of its members share a highly symmetrical TIM-barrel ((*β*/*α*)_8_) catalytic domain [Svensson, 1994] (Figure 1), including those structures without catalytic activity [Fort et al., 2007]. The TIM-barrel fold is highly versatile and widespread among the structurally characterized enzymes. It is present in almost 10% of all characterized enzymes [Farber, 1993, Höcker et al., 2001, Wierenga, 2001, Gerlt and Raushel, 2003].

**Figure 1:**
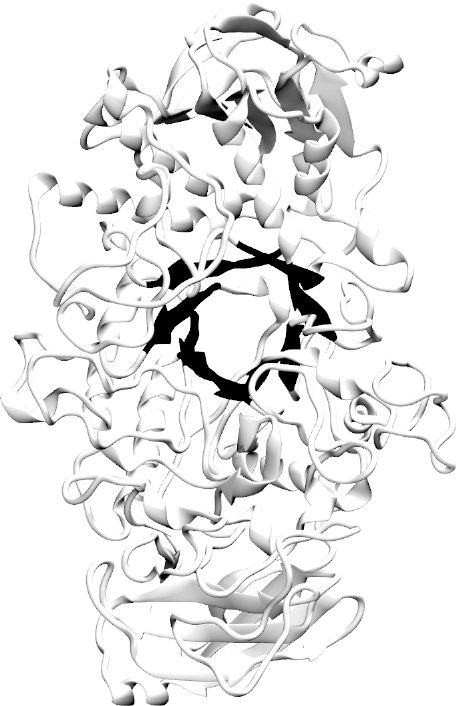
Structure of the catalytic domain of the *α*-Amylase. The TIM-barrel is highlighted. The image was rendered using VMD [Humphrey et al., 1996] and POVray (www.povray.org). The structure used to vizualize is the PDB 1BF2 chain A from *P. amyloderamosa*.

The GH13 TIM-barrel catalytic activity and substrate binding residues, occurs at the C-termini of *β*-strands and in loops that extend from these strands [Svensson, 1994]. The catalytic site includes aspartate as a catalytic nucleophile, glutamate as an acid/base, and a second aspartate for stabilization of the transition state [Uitdehaag et al., 1999]. The catalytic triad plus an arginine residue are conserved in this family across all catalysis-active members [Svensson and Janeček, 2015]. There are many well described enzymes in this family including: *α*-amylase (EC 3.2.1.1); oligo-1,6-glucosidase (EC 3.2.1.10); *α*-glucosidase (EC 3.2.1.20); pullulanase (EC 3.2.1.41); cyclomaltodextrinase (EC 3.2.1.54); maltotetraose-forming *α*-amylase (EC 3.2.1.60); isoamylase (EC 3.2.1.68); dextran glucosidase (EC 3.2.1.70); trehalose-6-phosphate hydrolase (EC 3.2.1.93); maltohexaose-forming *α*-Amylase (EC 3.2.1.98); maltotriose-forming *α*-Amylase (EC 3.2.1.116); maltogenic amylase (EC 3.2.1.133); neopullulanase (EC 3.2.1.135); malto-oligosyltrehalose trehalohydrolase (EC 3.2.1.141); limit dextrinase (EC 3.2.1.142); maltopentaose-forming *α*-Amylase (EC 3.2.1.-); amylosucrase (EC 2.4.1.4); sucrose phosphorylase (EC 2.4.1.7); branching enzyme (EC 2.4.1.18); cyclomaltodextrin glucanotransferase (CGTase) (EC 2.4.1.19); 4-*α*-glucanotransferase (EC 2.4.1.25); isomaltulose synthase (EC 5.4.99.11); and trehalose synthase (EC 5.4.99.16). It is a highly diverse family in both function and ubiquity, being found in all kingdoms of life [Svensson and Janeček, 2015]. Given sequence motifs and enzyme specificities [Cantarel et al., 2009] the GH13 family has been subdivided in over 40 subfamilies [Stam et al., 2006] all of which are closely related. The classification is based on the clustering of conserved regions [Janeček, 2002] and neighbour joining profiles [Stam et al., 2006], but they do not include the analysis of primary sourced structural information. Here we give insights into GH13 classification of subfamilies by analysing the family's structural variance on available solved structures.

## Material and Methods

### Homolog sampling

All available protein structures in the CAZY database that are classified as glycoside hydrolase family 13 (GH13) [Svensson and Janeček, 2015] were downloaded (Supplementary table S1). All metadata was also downloaded along with the PDB files.

A total of 388 structures were gathered (Supplementary table S1). Two protein structures had incomplete structural information (i.e. missing *C*_*α*_ in some residues). These structures were removed from the dataset for further analyses.

The remaining 386 structures where aligned using the algorithm proposed by Hleap et. al. [Hleap et al., 2013a] that modifies the pairwise MATT flexible structure aligner [Menke et al., 2008] to complete the multiple structure alignment of very large datasets. This procedure is performed because the multiple structural alignment version of MATT cannot reliable process such large dataset [Hleap et al., 2013a]. For this work, we used the quick heuristic search reported in Hleap et al. using 6 structures as starting points. To maximize the amount of homologous sites in the resulting alignment we used the Hamming distance as metric to select the best reference structure. Hleap et al. with Hamming distance as metric functions as follows:

1. Pick 6 random structures from the set (this will be increased by Hleap et al.
[Hleap et al., 2013a] algorithm to 18 starting structures) as reference structures.
2. Perform pair-wise alignments of every structure in the dataset to the reference structure.
3. Aggregate the alignments as a multiple structure alignment (MStA) having as reference the starting structures.
4. Re-code the MStA by homology information: 1 being homologous site, 0 non-homologous.
5. Evaluate each reference by the mean Hamming distance of to all other re-coded aligned structure’s sequence.
6. Pick the alignment that minimize the hamming distance, and thus maximizing the amount of structural data recovered.

### Geometrical analysis

#### Abstracting a protein structure as a shape

A protein fold can be defined as a 3D geometric shape. This is a novel application to protein structure of a well defined framework for bio-geometrical studies. Sequence analyses help to understand some trends, but explain little about geometry. Geometric Morphometric-like methods (GM) can be used to perform shape analysis from a geometric point of view. It also can be used to give insights into the phylogenetic relationships of the structures rather than the sequences. However, the application of GM to protein structures is not trivial. The scaling component of the Procrustes analysis have no conceptual equivalent for proteins. Since organisms grow it make sense to extract the size effect on shape in order to compare young with adults. On the other hand, in proteins the atoms do not stretch or grow, and therefore the scaling approach as proposed by [Adams and Naylor, 2000, 2003] is not relevant.

In the proposal of Adams and Naylor [2000, 2003], they: Abstract a residue as a landmark, evaluate its homology throughout the samples, using ClustalW [Thompson et al., 1994], delete gapped columns, and perform morphometric analyses.

The use of blind sequence alignment (a sequence alignment without structural information) to infer structural homology is not accurate, since the penalty for gaps in a loop region and the catalytic pocket may differ from other regions of the protein [Kann et al., 2005, Kjer et al., 2007]. Therefore, the definition of positional homology can differ. Since the structures are more conserved than their sequence, the alignment based on the structures have more reliable information of equivalent (homologous) residues across deeper phylogenetic sampling [Wohlers et al., 2012].

We used protein structural alignment strategies that have been studied extensively [Kolodny et al., 2005, Hasegawa and Holm, 2009, Poleksic, 2011, Joseph et al., 2011, Shibberu et al., 2012]. Here, the Multiple Alignment with Translations and Twists (MATT) algorithm [Menke et al., 2008] was used. This approach strips out rotational and translational information, as well as the variability induced by flexible hinges along the backbone.

The abstraction of the residues and landmarks is similar to that in Adams & Naylor [Adams and Naylor, 2000, 2003]. Here we assign a landmark to residue centroids defined by (x,y,z):

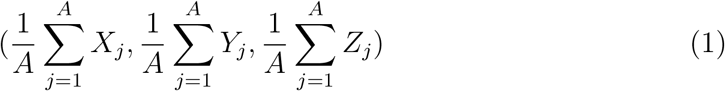

where *A* will be the number of heavy atoms that constitutes the side chain of a residue including the *C*_*α*_. This procedure takes into account only the strictly homologous residues (no gaps in the alignment). It includes the variance of both the backbone, and side chain. In the case of glycine the centroid is the *C*_*α*_ itself.

#### Structural clustering

To analyse the structural similarity of all proteins, a clustering on a graph abstraction is used. Let *S* = (*N, f*) be a fully connected undirected graph representing the set protein structures with *N* being the individual structures abstracted as nodes. *f* will be a function *f*: *N* × *N* → 𝕜 that assigns the weight to each edge. The weight here is defined as 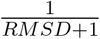 between the two nodes. To avoid biases towards a reference we used the pairwise version of MATT to compute all pairwise alignments and obtain their similarity measures. After defining the graph, the community structure is assessed using a fast-greedy approach to maximize the modularity index (*Q*) as reported in Newman [Newman, 2004]. The graph construction and clustering was performed with the python module for igraph [Csardi and Nepusz, 2006].

The clusters are further tested for statistical significance following the method reported in Hleap et al. [Hleap et al., 2013b].

After a stable clustering scheme has been found, a hierarchical clustering is then tested. Each cluster found is re-clustered and tested for significance using the same approach until no further clusters are produced. This approach will explore subclusters that may be meaningful in protein structure classification.

#### Analysis of the deformation in superimposed structures/shapes

The inter-landmark distance matrix (form configuration) is computed using the Euclidean distance for each entry in *m* dimensions:

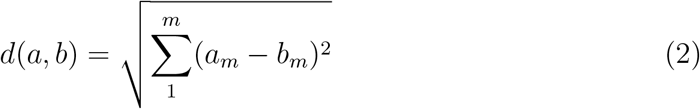

where *d*(*a*, *b*) stands for the Euclidean distance between variables *a* and *b*. Therefore the form matrix (*FM*) is [Claude, 2008]:

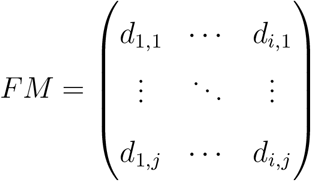

*FM* is then a square symmetric matrix, with zeros in the diagonal entries. If two forms (shapes in the inter-landmark framework) are identical, they will have the same entries in the *FM* matrix. The matrix of differences in form (The form difference matrix or FDM as named by Claude [Claude, 2008] between two configurations *S*1 and *S*2 is given by:

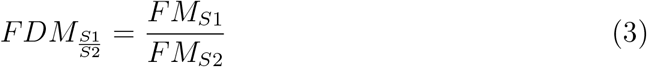

The score of the most influential point (*I*) in the data can be computed by adding the sum of the differences to the median value per column (variable) and ranking the positions. This can be computed as:

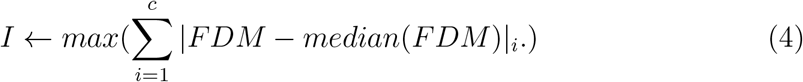

where *c* is the column index, and FDM is the form difference matrix.

However, as shown in equation 3, this FDM is the representation of the difference between two shapes. We can generalize this by summing up the residuals of all shapes versus an hypothetical mean shape. For simplicity this can be calculated as the per-variable per-dimension average. In other words, the average of each dimension of each landmark. This approach will then return a Form Difference (FD) value per landmark, however; this value is not bounded and is difficult to interpret. For this reason we scaled the resulting FD vector 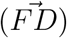 such that it is bounded from −1 (least variation) to 1 (highest variation) with:

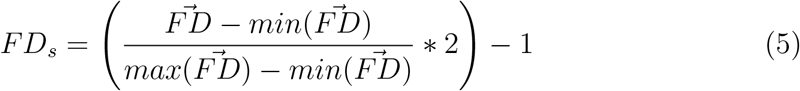

This analysis is performed in each group of aligned structures. The groups can be any level of the clustering that was explored in the previous section.

All statistical analysis and protein structure manipulation were performed with R and python scripts.

### Phylogenetic analysis

The phylogenetic tree was constructed using RAxML v.8 [Stamatakis, 2014], using the output of FastTree [Price et al., 2010] as starting tree. The best protein substitution model was automatically determined by RAxML. The extended majority rules method (MRE) bootstopping criteria [Pattengale et al., 2009] was used to infer the robustness of the tree with the appropriate bootstrap replicates. The visualization of the tree was performed with FigTree v1. 3.1 [Rambaut, 2009].

## Results & Discussion

### Structural groups

Three major structural clusters where found, based on protein structure similarity:

#### Animal-like *α*-Amylases

One hundred and nine structures were classified in this group based on their RMSD similarity. Animal *alpha*-amylases are allosterically activated by chloride ions [D’Amico et al., 2000]. Chloride ion binding in animal amylases is performed by an arginine, an asparagine, and another basic residue. This binding site is located within 5 Å of the active site and the substrate binding cleft facilitating the catalytic activity [Maurus et al., 2005]. The chloride dependency can be found also in the psychrophilic bacterium *Pseudoalteromonas haloplanktis*. In the bacterial *α*-amylase, the chloride-binding arginine is replaced by lysine. D’amico et al. [D’amico et al., 2000] and DaLage et al. [Da Lage et al., 2004] demonstrated a high similarity between animal amylases and the *P. haloplanktis α*-amylase. These authors proposed that the similarity is explained by a horizontal gene transfer between animals and bacteria [Da Lage et al., 2004]. Given the phylogenetic and structural proximity between animal *P. haloplanktis* amylases (Figure 2), we provide further evidence for such an event. Despite that, the mechanisms to which an Antarctic bacterium obtained an animal gene remains elusive. A plausible hypothesis would involved a marine arthropod, with an *α*-Amylase more closely related to the insect *α*-amylases (Figure 2).

**Figure 2:**
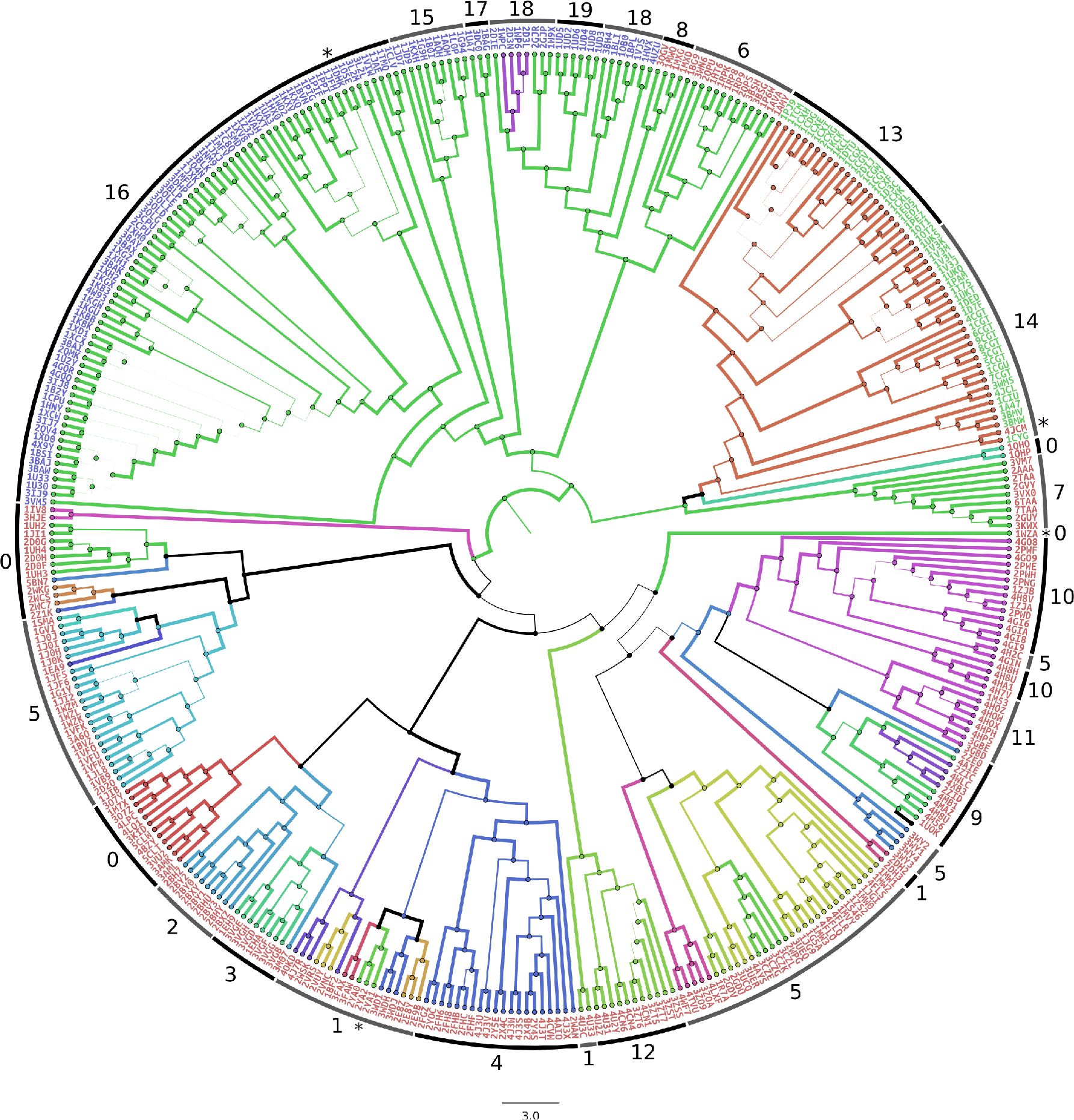
Maximum likelihood tree of the *α*-Amylase family. Branch thickness corresponds to its bootstrap value. Branch color corresponds to its enzyme function codification (EC). Taxa labels correspond to the PDB codes and they are colored based on the top level clustering. Numbers above the taxa labels are the numerical representations of the last level of hierarchical clustering.

Another group of bacterial amylases were recovered as part of the animal-like *α*-amylases, stressing their structural similarity. Interestingly, the bacterial and archaeal amylases are composed of structures that either have been engineered to increase stability and catalytic efficiency (e.g. 4UZU; [Offen et al., 2015]) or are extremophiles with highly efficient *α*-Amylases (e.g 1W9X; [Davies et al., 2005]).

Structurally, this is a more or less conserved group and is also very functionally homogeneous. A small clade within the *Bacillus* genus with an Glucan 1,4-alpha-maltohexaosidase (or G6-amylase; EC 3.2.1.98) instead of the *α*-amylase (EC 3.2.1.1) activity can be seen from Figure 2. However, all these bacteria are alkalophilic: They are very stable and use different substrates than their non-alkalophilic counterparts.

When the hierarchical clustering is applied to this group, it can be broken down into 5 subclusters (Figure 2): *P. haloplanktis α*-amylases (Cluster 15), Animal *α*-amylases (Cluster 16), engineered *Bacillus alpha*-amylases (Cluster 17), bacterial *α*-amylase (Cluster 18), calcium-free *α*-amylase (Cluster 19). Cluster 15 is a non-monophyletic group consisting in the Arthropod/*Pseudoalteromonas α*-Amylases. This cluster is split into subfamilies 15 and 32 according to the CAZY classification [Stam et al., 2006]. The aggregation of these two families under the sub cluster 15 makes sense in the light of the ancient horizontal gene transfer (HGT) hypothesis proposed by DaLage et al [Da Lage et al., 2004]. If the ancient HGT happened between a marine arthropod and a ancestor of the bacterium, the structural similarity should be evident. Here we show that despite the phylogenetic disparity between the insect and *Pseudoalteromonas*, the structurally similarity suggests an HGT between a marine arthropod and a marine bacterium. This observation is not evident from the CAZY subfamily classification.

The deformation analysis based on FD (see methods) was performed in a secondary alignment of the 109 structures. This alignment was optimized to recover more data by removing structural outliers such as smaller structures. The resulting alignment contained 85 structures corresponding to the animal-like *α*-amylases (Clusters 15 and 16; Figure 3).

**Figure 3:**
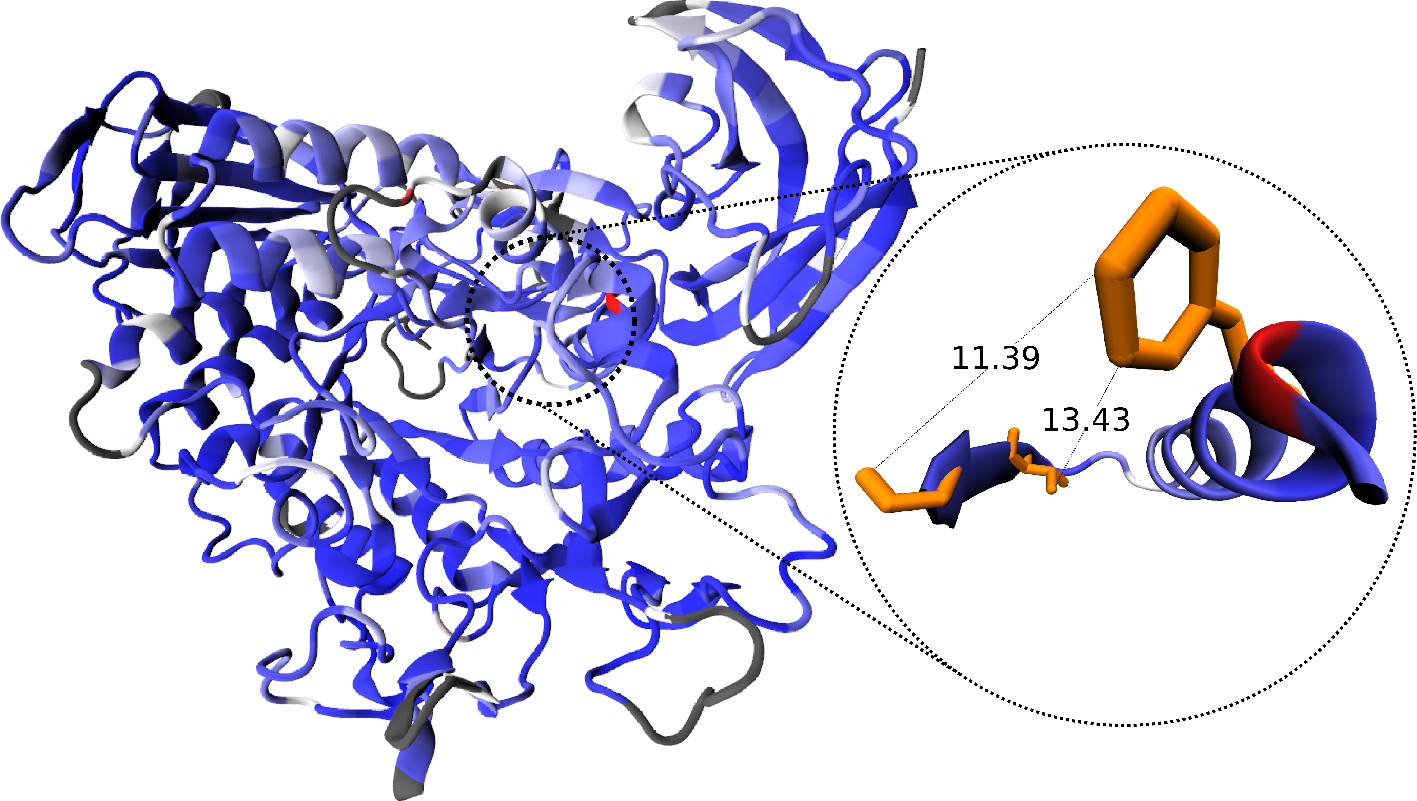
Scaled form difference for the animal-like *α*-Amylase cluster. A) Surface representation of the porcine *α*-amylase (1PPI), colored by FD. B) The most conserved (lowest FDs) hubs. The form difference scaled values rendered into the reference structure 1PPI from *Sus Scrofa*. The blue to red coloring represents the scaled FD, ranging from -1 (blue) to +1 (red). Lycorice orange resiudes correspond to the caralytic triad.

The FD analysis shows a tendency of this subgroup to possess structurally conserved cores despite some sequence variation (Figure 3A; Supplementary material S1) that builds up to a more movable surface (Figure 3B, Supplementary material S1). It is interesting, however, that the catalytic pocket is not the most geometrically conserved, showing a higher degree of flexibility than its scaffold. This observation might be related to intrinsic flexibility to the uptake and catalysis of carbohydrates, including some small degree of conformational change. The least variable residues appear to be near the surface and correspond to 135D and 174Y in the structure 1PPI. These two positions are highly conserved both in sequence and geometry, and seem to hold together the two sides of the TIM-barrel by a salt bridge. The positions 372I, 373N, 377T, and 405P appear to be the major contributors to the shape deformation. Interestingly, all these positions are highly conserved in this group. The former three residues are in a cluster after the 378C-384C bridge, and their contribution to the deformation might be due to the lack of of charge and the lack of neighbouring serines. This feature might be to stabilize and maintain the loop created by the Cysteine bridge. In the case of the Proline, its contribution to deformation might be due to the lack of interactions with its neighbours which gives the constitution of loop to this region. In this case, as before, the conservation of this secondary structure might help to the transition from a mainly alpha helical structure to *β*-sandwich, and therefore helping to keep the structural consistency of the TIM-barrel.

The remaining 24 structures were also aligned and an FD analysis performed. All these 24 structures correspond to the animal-like bacterial *α*-amylases. The analysis show a similar trend that in the animal *α*-amylases, however, the catalytic pocket and the TIM-barrel showed more rigidity. The most structurally conserved sites seem to be clustered in the domain B (as referred by Offen et al. [Offen et al., 2015]) situated in between the *β*-strand 3 and the *α*-helix 3 (See Supplementary material S2).

#### Cyclodextrin glycosyltransferase group

This structural group is the most geometrically conserved of the recovered clusters. Cyclodextrin glycosyltransferases (EC 2.4.1.19; CGTase) are extracellular enzymes with capability of forming cyclodextrins from starch and mainly found in bacteria [Xie et al., 2014]. They can produce *α*, *β* and *γ*-cyclodextrins (with 6, 7, or 8 glucose residues), or even larger products [Leemhuis et al., 2002]. In the CAZY database, this group is classified as subfamily 2 of the GH13 family. In Figure 2, this group is split into two (clusters 13 and 14). Interestingly, this split relates to the prevalence of *α* (cluster 14) or *β* (cluster 13) cyclodextrins. That is, our method was able to identify subtle changes in the geometry of the proteins to detect preferences for larger or smaller products. The cluster 14 is polyphyletic with respect to one member (4JCM; asterisk within cluster 14 in Figure 2). This structure shows a higher prevalence for *γ*-cyclodextrins, and therefore for larger products. The structural shift in this structure made it more structurally related to cluster 0 than cluster 14. This is an interesting observation since by sequence-based clustering 4JCM belongs to the same group as the the rest of structures belonging to cluster 14. This means that the methods employed here are sensitive to structural shifts, and might be of interest for further classification or re-classification of the CGTases.

The statistical analysis on the deformation of this set of proteins is rendered in figure 4. The most deforming element was found to be Gly659 (3WMS numbering), which was found in a loop as expected. Interestingly, the second most deforming residue in the structure the highly conserved Phe236 (3WMS numbering) that is located within 15 Å of the catalytic residues. It seems that this residue is important for the uptake of the ligand or is related to the size of the product that would be released.

**Figure 4:**
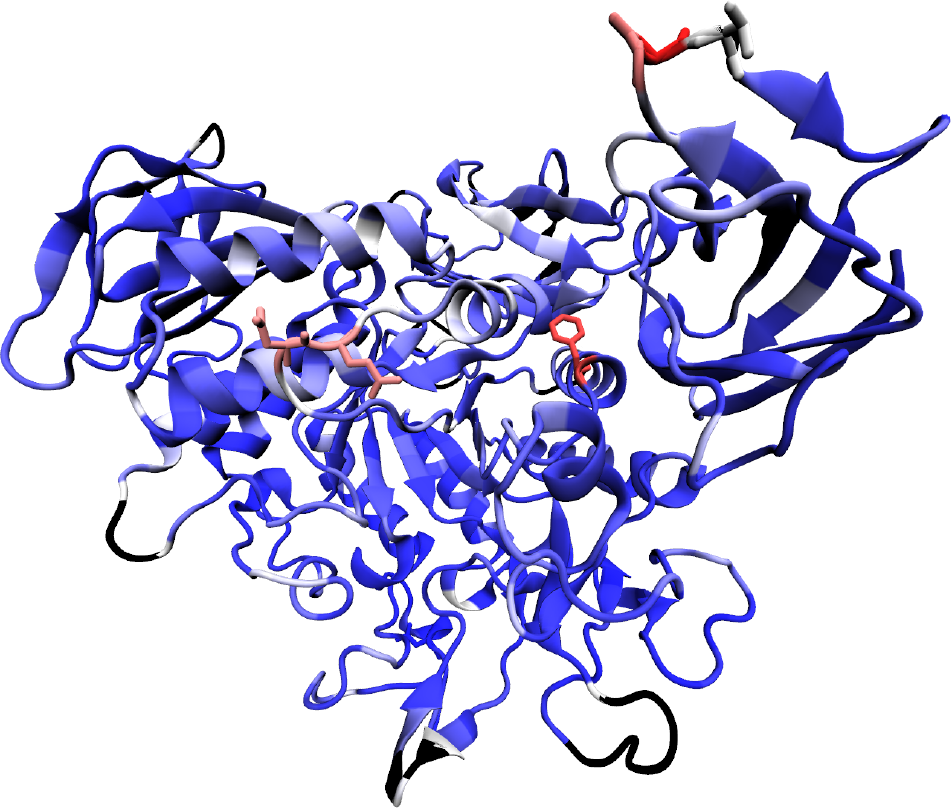
Scaled form difference for the bacterial cyclodextrin glucanotransferases cluster. The form difference scaled values rendered into the reference structure 3WMS from *Paenibacillus macerans*. The blue to red coloring represents the scaled FD, ranging from −1 (blue) to +1 (red). Lycorice resiudes correspond to the highest FD.

By the nature of the clustering algorithm, some of the outliers tend to cluster together to the exclusion of the most cohesive groups. In our results, cluster 0 contain more structures than the other higher level clusters. It is also the most structurally and functionally diverse. In the hierarchical clustering was further divided into 11 subclusters with an apparent high structural and functional diversity. This group should be further explored to explained the diversity found, however, with the current experimental information available is not possible to give support to the diversity observed. Nevertheless, it poses interesting structural questions into the classification and evolution of the GH13 family. We have observed that when tight clusters are present amongst more variable samples, often those more variable samples tend to group into a cluster at the top level of hierarchy.

## Conclusions

We have provided evidence supporting the grouping of the GH13 families. We also provided evidence for further subdivision (e.g. cluster 13 and 14), based solely on structural information. This new information provides interesting basis to refine or modify the current classification, as well as the selection of candidates for engineering studies. Furthermore, we provided evidence supporting the HGT hypothesis between a bacterium and an animal. We posit here that the most plausible scenario happened between an ancestral arthropod and an ancestral (probably gut associated) bacterium. We also introduced the form difference analysis to identify potential candidates for mutagenesis and possible functional and evolutionary analyses. Despite the utility of the later, its interpretation should be always be with respect to the given dataset, and more work is needed in order to be comparable among studies. One possibility to make the scaled FD more comparable is to use it as coefficient of the entropy of each site, in such a way that the relative importance of the residue can be highlighted. By including degree of information in each site (entropy) and weighted by the geometrical conservation (FD) the relative importance of each site might be more accurate. This is due to the fact that a low information site (high entropy) is expected to have a higher geometrical deformation and vice-versa. Any deviations from these expectations provide information into structurally important sites. It is therefore important to analyse these sites, and develop this method further.

## Acknowledgements

The authors thank the members of the Blouin Lab for helpful comments and critical review of this manuscript. This study was funded by NSERC through the grant No. 120504858. This work was partially supported by the Departamento Administrativo de Ciencia y Tecnología - Colciencias (Colombia) through the CALDAS scholarship.

